# PI31 expression prevents neuronal degeneration in a mouse Parkinson Disease model

**DOI:** 10.1101/2020.05.05.078832

**Authors:** Adi Minis, Hermann Steller

## Abstract

Age-related neurodegenerative diseases pose a major unmet health need since no effective treatment strategies are currently available. These disorders are defined by the accumulation of abnormal protein aggregates that impair synaptic function and cause progressive neuronal degeneration. Therefore, stimulating protein clearance mechanisms may be neuro-protective. The proteasome regulator PI31 promotes local protein degradation at synapses by mediating fast proteasome transport in neurites, and loss of PI31 function causes neuronal degeneration. Here we show that transgenic expression of PI31 in a mouse Parkinson’s Disease model preserves neuronal function and greatly extends animal health and lifespan. These results indicate that targeting the PI31-pathway may have therapeutic value for treating neurodegenerative disorders.

---

A common feature of neurodegenerative diseases, such as Alzheimer’s Disease (AD), Parkinson’s Disease (PD), and Amyotrophic Lateral Sclerosis (ALS) is the accumulation of abnormal protein aggregates ^1–3^. However, the precise mechanisms underlying the accumulation of pathognomonic proteins and the nature of their toxicity have remained elusive^4–7^. Although these conditions affect different types of neurons, they all begin with alterations in synaptic efficacy that are associated with synaptic and dendritic spine pathology, pointing to synaptic failure as a common and early disease mechanism^8,9^. Moreover, the various pathological aggregates and deposits in these diseases contain poly-ubiquitinated (poly-Ub) proteins, indicating that these proteins were tagged for destruction but escaped proteasome-mediated degradation^10^. One possible explanation is that insufficient local activity of proteasomes at synapses initiates the formation of poly-Ub aggregates. The ubiquitin-proteasome system (UPS) is the primary mechanism by which cells degrade unwanted, damaged and potentially toxic intracellular proteins^4,11^. Local protein degradation at synapses is critical for synaptic function and plasticity^12–14^. This requires long-distance transport of proteasomes between the neuronal cell bodies and synapses in the periphery. The conserved proteasome-binding protein PI31 mediates fast transport of proteasomes in axons and dendrites, and disruption of this process impairs synaptic protein homeostasis and causes progressive neuronal dysfunction and degeneration^15,16^. The PI31-pathway is linked to human disease because mutations in both PI31 and its conserved binding partner, Fbxo7/PARK15 have been identified in human patients^17–21^. Mutations in Fbxo7/PARK15 cause Parkinsonian Pyramidal Syndrome in humans and Parkinsonian-like phenotypes in mice^22,23^. Significantly, inactivation of Fbxo7/PARK15 leads to proteolytic cleavage and reduced levels of PI3 (Fig. 1A)^22,24^. This suggests that loss of Fbxo7/PARK15 may cause disease by attenuating PI31 function.

**Fig. 1.**
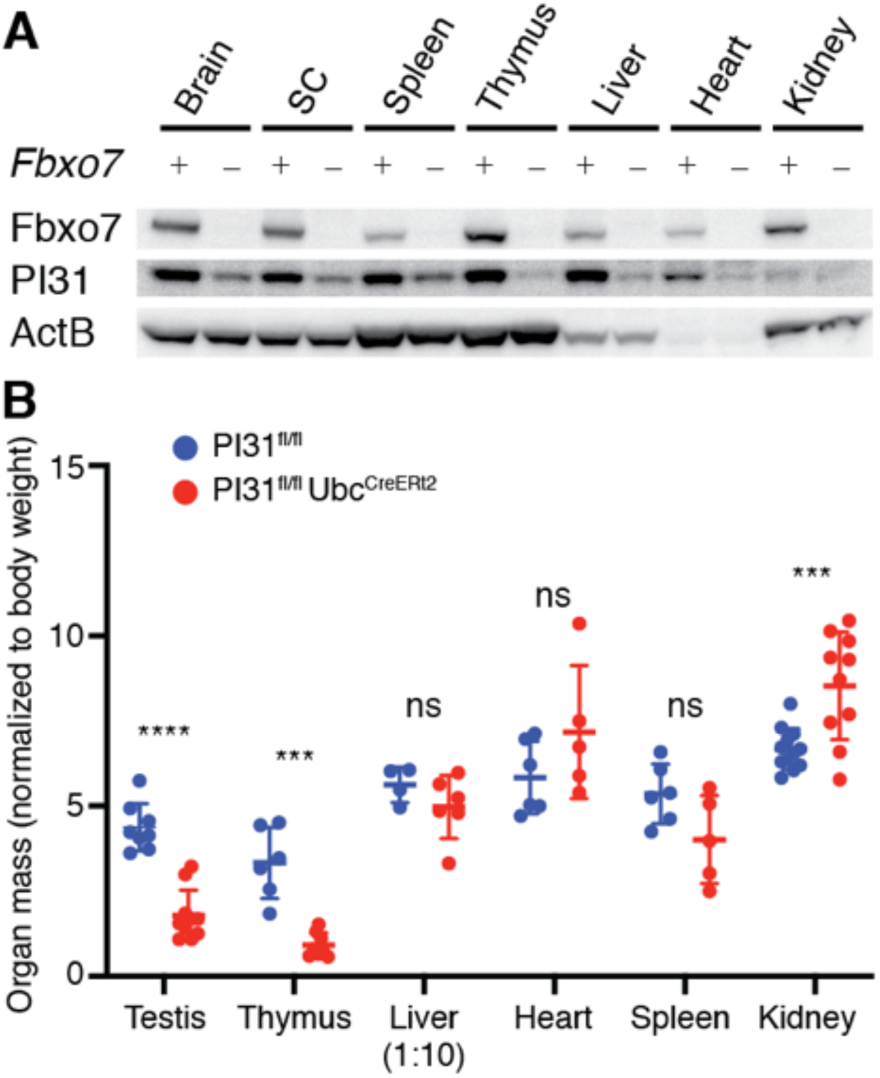
Inactivation of Fbxo7 reduces PI31 protein levels. (A) Western blot analysis revealed that loss of *Fbxo7* reduces PI31 protein levels in different tissues. Control n=3, Fbxo7 knockout n=3 (B) Conditional inactivation of PI31 in adult mice leads to a dramatic reduction in thymus and testis size and weight. Control n=4-6, Fbxo7 knockout n=5-8. Statistical significance was defined by student t-test. Data is represented by a grouped scatter plot with mean and SD. *** pValue < 0.001, **** pValue <0.0001, ns = non significant.

To investigate this idea, we first compared phenotypes resulting from conditional inactivation (CKO) of PI31 to those of Fbxo7 and found them to be remarkably similar. Like Fbxo7 mutant mice, PI31 CKO mice have a dramatic decrease in testis and thymus size with similar histopathology (Fig. 1B, S1)^25^. Moreover, inactivation of Fbxo7 in mouse motor neurons produced phenotypes strikingly similar to PI31 CKO (Fig. 2A)^16^. Like for PI31^fl/fl^;Hb9^Cre^, Fbxo7^fl/fl^;Hb9^Cre^ mice were born with the expected Mendelian ratio and initially showed no gross motor dysfunction. Four weeks after birth, a difference in weight, kyphosis and motor problems became apparent that progressively increased with age (Fig. 2A and data not shown). We used whole-mount fluorescent confocal imaging to examine motor innervation of the Triangularis Sterni muscle and saw a loss of synapse architecture, axonal swellings, axon sprouting and large P62 granules in the NMJ and its vicinity, very similar to our PI31 CKO (Fig 2B-H)^16,26^. Therefore, motor neuron-specific loss of either Fbxo7 of PI31 produces very similar phenotypes. If the root cause of neuronal pathology seen in Fbxo7^fl/fl^;Hb9^Cre^ is through decreased PI31 function, ectopic expression of PI31 should be able to rescue Fbxo7 deficiency. To test this hypothesis, we generated transgenic mice that conditionally expresses a Flag-tagged form of PI31 (TgPI31) (Fig. 3A). Since the Hb9^Cre^ allele is not viable as a homozygous, we used the Chat^Cre^ driver for all experiments from here on to increase the number of animals with desired genotypes. Chat^Cre^ has a broader expression pattern than Hb9^Cre^, which includes spinal motor neurons along with all other cholinergic neurons. As expected, both physiological and cellular phenotypes were more severe compared to Hb9^Cre^. In order to validate that TgPI31 is expressed and functional, we tested its ability to rescue PI31 mutant mice (Fig. 3B-H). The TgPI31 construct expressed protein of expected size and was able to completely rescue PI31^fl/fl^;Hb9^Cre^ mutant mice, demonstrating that the transgenic protein is biologically active (Fig. 3B-H). Next, we crossed TgPI31 to Fbxo7^fl/fl^;Chat^Cre^ mice to test for possible rescue. Fbxo7^fl/fl^;Chat^Cre^ were born at expected Mendelian ratios and appeared similar to their control littermates in the first 3 weeks of age (Movie S1). However, starting at 4 weeks of age, mutant mice failed to gain weight and by the age of 8 weeks had to be euthanized due to poor health (Fig. 4A,B). Again, the phenotypes of Fbxo7^fl/fl^;Chat^Cre^ were very similar to PI31^fl/fl^;Chat^Cre^ CKO at both the cellular and behavioral level. In particular, the synaptic architecture of the NMJ in Fbxo7^fl/fl^;Chat^Cre^ mice was highly abnormal, and neurons showed pathological hallmarks, such as axonal swelling and sprouting (Fig. 4C,D,F,G,I and J).

**Fig. 2.**
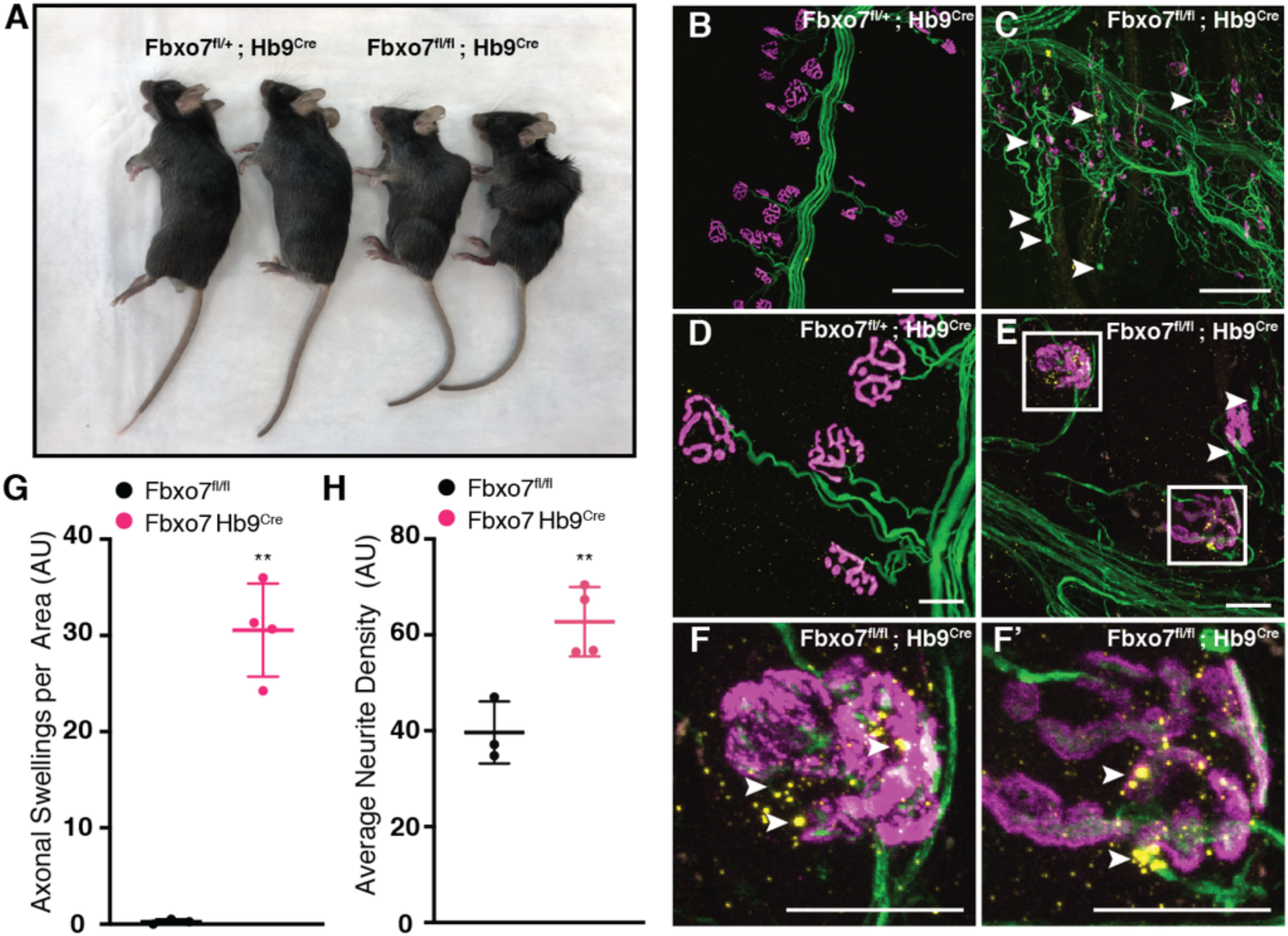
Motor neuron-specific inactivation of Fbxo7 resembles loss of PI31 function. (A) 3-month old Fbxo7^fl/fl^;Hb9^Cre^ and Fbxo7^fl/fl^ control mice. Fbxo7^fl/fl^;Hb9^Cre^ mice lost weight and developed kyphosis, similar to PI31^fl/fl^;Hb9^Cre^ mice^16^ (B-F) Micrographs depict the innervation of Triangularis Sterni muscles by spinal motor neurons. Loss of Fbxo7 resulted in axonal swellings (white arrowheads), especially at synapses and their vicinity, and axonal sprouting (B-E). Accumulation of large P62 aggregates at NMJs of Fbxo7^fl/fl^;Hb9^Cre^ mice is indicative of proteotoxic stress (F, F’ are insets marked by white boxes in E). (G,H) Quantification of axonal swellings and axon sprouting. Scale bars in B,C 100μm, in D-F 20μm. Statistical significance was defined by student t-test. Data is represented by a grouped scatter plot with mean and SD. ** pValue < 0.01. Control n=3, Fbxo7^fl/fl^;Hb9^Cre^ n=4.

**Fig. 3.**
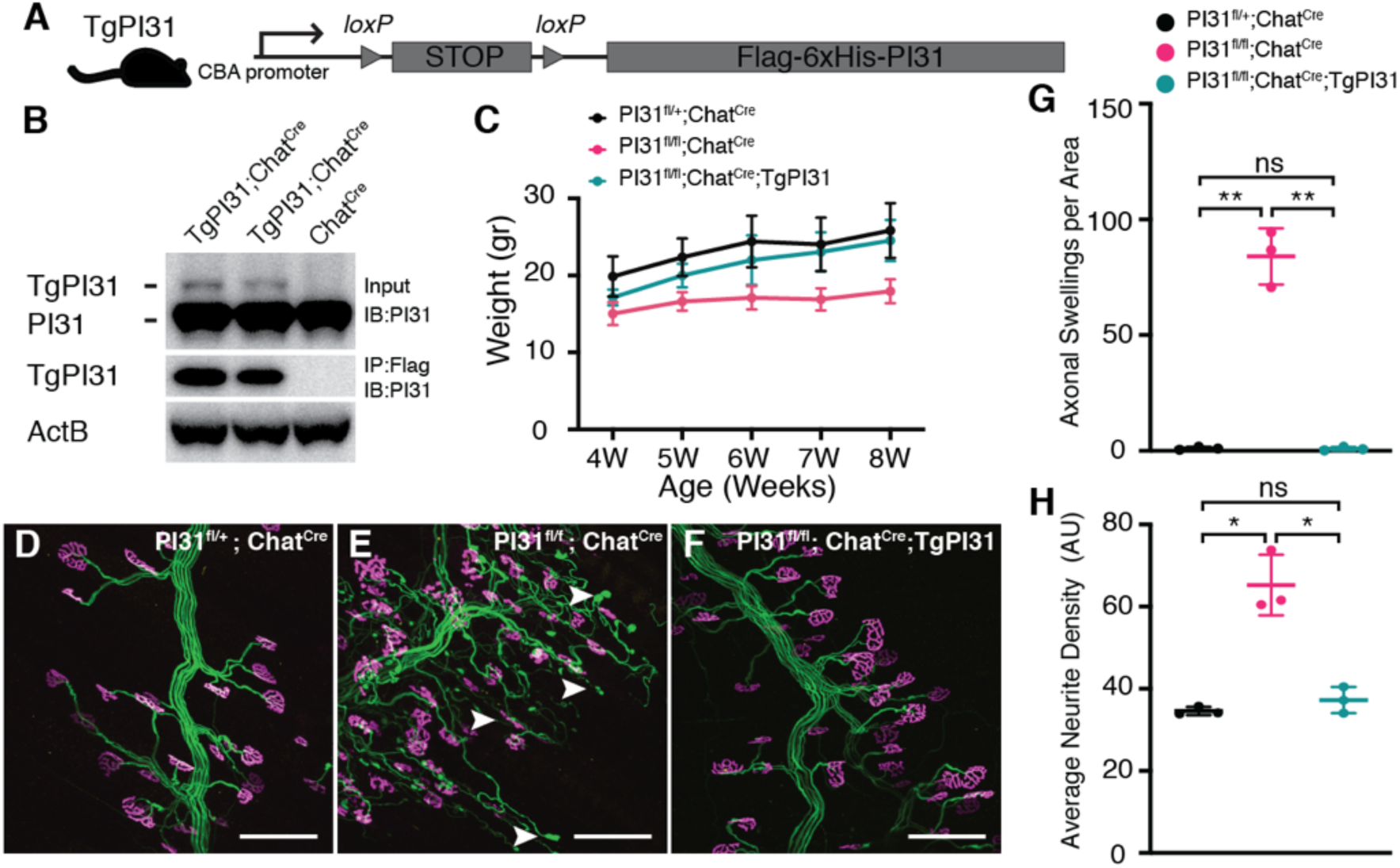
Conditional expression of PI31 in transgenic mice. (A) Design of the TgPI31 transgene. A floxed STOP cassette was inserted upstream of a flag-his tagged PI31 open reading frame to allows expression of PI31 only after cre-mediated recombination. (B) Immunoprecipitation and Western blot analysis of spinal cord protein extracts was used to demonstrate proper inducible expression of ectopic PI31 in TgPI31 Chat^Cre^ mice. The upper band in the TgPI31;Chat^Cre^ input lanes represents the expected size of the tagged transgenic PI31 protein. Immunopreciptation was done using anti-Flag tag magnetic beads. Blots were done with anti-PI31 antibody. Presented are two TgPI31;Chat^Cre^ biological replicates. (C-H) Cholinergic neuron-specific loss of PI31 can be rescued by transgenic expression of recombinant PI31, demonstrating that the tagged PI31 protein is biologically active. (C) Mouse body weight of PI31, control mice, PI31^fl/fl^;Chat^Cre^ and PI31^fl/fl^;Chat^Cre^;TgPI31 mice. PI31^fl/fl^;Chat^Cre^ had reduced weight, but transgenic expression of PI31 (PI31^fl/fl^;Chat^Cre^;TgPI31) restored body weight to that of control littermates. (D-F) Micrographs of Triangularis Sterni muscle innervation by spinal motor neurons. Loss of PI31 in cholinergic neurons resulted in axonal swellings (white arrowheads) and axonal sprouting, but transgenic expression of PI31 rescued these phenotypes. (G,H) quantification of axonal swellings and axon sprouting. Scale bars in D-F 100μm. Statistical significance was defined by student t-test. Statistical significance was defined by student t-test. Data is represented by a grouped scatter plot with mean and SD. * pValue < 0.05, ** pValue < 0.01. Control n=3, PI31^fl/fl^;Chat^Cre^ n=3, PI31^fl/fl^;TgPI31;Chat^Cre^ n=3.

**Fig. 4.**
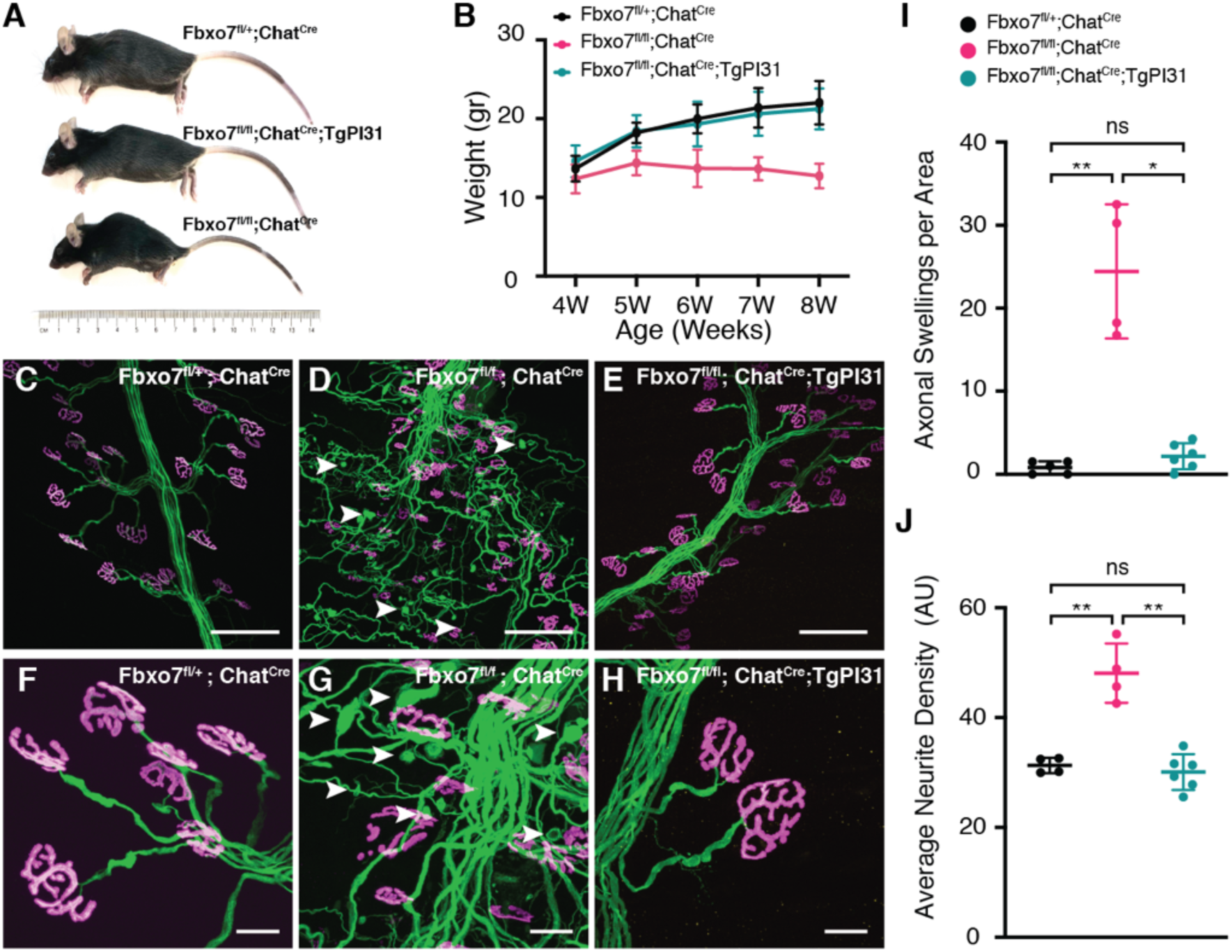
Transgenic expression of PI31 rescues Fbxo7 knockout pathology. (A) 8-week old Fbxo7^fl/fl^;Chat^Cre^, Fbxo7 control and Fbxo7^fl/fl^;Chat^Cre^;TgPI31. Fbxo7^fl/fl^;Chat^Cre^ animals were significantly smaller in size and developed kyphosis. Transgenic expression of PI31 in these mice (Fbxo7^fl/fl^;Chat^Cre^;TgPI31) suppressed these phenotypes and yielded animals that were indistinguishable from normal control littermates. (B) Quantification of mouse body weight. Fbxo7^fl/fl^;Chat^Cre^ loose weight, transgenic expression of PI31 in Fbxo7^fl/fl^;Chat^Cre^;TgPI31 mice restored body weight to that of control littermates. (C-H) Micrographs of Triangularis Sterni muscle innervation by spinal motor neurons. Loss of Fbxo7 in cholinergic neurons results in axonal swellings (white arrowheads) and axonal sprouting. These phenotypes were rescued to control levels in mice that ectopically express TgPI31. (I,J) quantification of axonal swellings and axon sprouting. Scale bars in C-E 100μm, in F-H 20μ. Statistical significance was defined by student t-test. Data is represented by a grouped scatter plot with mean and SD. * pValue < 0.05, ** pValue < 0.01. Control n=4, Fbxo7^fl/fl^;Chat^Cre^ n=4, Fbxo7^fl/fl^;TgPI31;Chat^Cre^ n=6.

Strikingly, whereas Fbxo7^fl/fl^;Chat^Cre^ developed a progressive motor neuron disease, Fbxo7^fl/fl^;TgPI31;Chat^Cre^ mice were practically indistinguishable from control animals and did not develop any detectable motor dysfunction (Fig. 4A,B, Movie S2). Micrographs of muscle innervation confirmed that NMJs of transgenically rescued animals were intact and appeared similar to that of normal controls (Fig. 4E, H, I and J). We conclude that expression of PI31 in Fbxo7 CKO motor neurons can prevent axonal degeneration and preserve neuronal function and organismal health. This indicates that neuronal degeneration caused by mutations in Fbxo7/PARK15 is caused, to a significant extent, through attenuation of PI31 function.

Our findings conflict with an earlier report on Fbxo7 CKO mice^22^. Although this study also showed that loss of Fbxo7 reduces PI31 protein levels, the authors argued that proteasomal defects were not mediated by PI31^22^. However, their conclusions were based on cell-wide assays for proteasome activity, which would have missed localized defects due to impaired proteasome transport. We do not wish to rule out that Fbxo7/PARK15 plays other important roles beyond promoting PI31 stability, as previously suggested^27,28^. Yet, our ability to extensively attenuate Fbxo7 mutant phenotypes in both *Drosophila* spermatids and mouse neurons indicates that PI31 is a very important biological target for Fbxo7, and that this pathway has been conserved in evolution from insects to mammals^24^. Our findings also have important implications for the possible development of therapeutic strategies to treat age-related neurodegenerative diseases associated with aggregate-prone proteins. Impaired activity of the UPS is well documented in age-related neuronal degeneration, and efforts to stimulate proteasome activity to promote clearance of aggregate-prone pathognomonic proteins have been made^29,30^. However, increasing the local proteolytic capacity at synapses through targeting proteasome transport pathways has not been previously explored as the required tools were lacking. The results presented here suggest that targeting the PI31 pathway is a promising strategy in this regard.

## Materials and Methods

### Mice

All animal work was performed as required by the United States Animal Welfare Act and the National Institutes of Health’s policy to ensure proper care and use of laboratory animals for research, and under established guidelines and supervision by the Institutional Animal Care and Use Committee (IACUC) of The Rockefeller University. Protocol was approved by Dr. Engin Ozertugrul PhD M.Ed. Mice were housed in accredited facilities of the Association for Assessment of Laboratory Animal Care (AALAC) in accordance with the National Institutes of Health guidelines.

### Generation of Fbxo7 transgenic mice

Fbxo7<tm1a(EUCOMM)Wtsi>/WtsiCnrm) cryo-preserved sperm was purchased from the European Mouse Mutant Archive (EMMA – INFRAFRONTIER). EMMA protocol for In Vitro Fertilization was followed to generate the Fbxo7 tm1a ‘Knockout First’ mouse strain. Fbxo7 tm1a heterozygous mice were then bred to CMV^Cre^ (Jackson Laboratories) mice or Actin^Flp^ (Jackson Laboratories) in order to generate heterozygous Fbxo7 Null (tm1b) and Fbxo7 conditional (tm1c) alleles, respectively.

### Generation of TgPI31 transgenic mice

The mouse PI31 gene was amplified by PCR using primers listed below. NheI and KpnI restriction sites were added to the forward and reverse primers, respectively. Flag and 6xHIS tags sequenced were also added to the 5’ primer. PCR products were purified from agarose gels and digested using NheI and KpnI restriction enzymes. Flag-6xHIS-PI31 was cloned into pCAAGS\ES using NheI and KpnI RE cloning. To generate a conditional construct, Flag-6xHIS-PI31 was inserted downstream of a floxed “STOP” cassette using EcoRI and NheI RE cloning into the pB-RAGE plasmid. The plasmid was then digested using SalI and Eco147I (StuI) RE. DNA fragment purification and pronuclear injection for the generation of transgenic mice was preformed by the Transgenic and Reproductive Technology Center of the Rockefeller University. Pups were genotyped and transgene constructs were sequenced to ensure proper incorporation into host DNA.

Generation of PI31 conditional and inducible transgenic mice was described previously (16).

In order to generate motor neuron and cholinergic neuron conditional strains, we bred mice to Hb9^Cre^ (Jackson laboratories) and Chat^Cre^ (Jackson laboratories) strains respectively.

### Genotyping

Tails were lysed in 150μl of direct tail lysis solution supplemented with proteinase K, and incubated for 4-8 hours at 55 C. Proteinase K was heat-inactivated at 90c for 30 minutes. PCR was performed with OneTaq PCR polymerase. 1ul tail suspension was used as a PCR template. Genotyping PCR program

Genotyping PCR program
| Step | Temp C | Time | Note |
| --- | --- | --- | --- |
| 1 | 94 | 2' |  |
| 2 | 94 | 20'' |  |
| 3 | 65 | 15'' | Decrease -0.5c per cycle |
| 4 | 68 | 1' | Repeat steps 2-4 for 10 cycles (touchdown) |
| 5 | 94 | 15'' |  |
| 6 | 55 | 15'' |  |
| 7 | 72 | 1' | Repeat steps 5-7 for 28 cycles |
| 8 | 72 | 2' |  |
| 9 | 10 | Hold |  |

Expected amplicon size:

| Name | Size (bp) |
| --- | --- |
| PI31 wt | 250 |
| PI31 floxed | 400 |
| Fbxo7 wt | 800 |
| Fbxo7 floxed | 600 |
| Fbxo7 ko (LacZ) | 400 |
| TgPI31 | 350 |
| Cre | 250 |
| Flpe | 100 |

### Tamoxifen Injection

For Tamoxifen injections, tamoxifen was dissolved in corn oil over night at 37 C to a final concentration of 20mg/ml. 100μl of tamoxifen solution was injected IP into 6-week old mice for 5 consecutive days. Mice were sacrificed for necropsy two and a half weeks after the last injection was administered.

### Western Blots and Immuno-precipitation

Fbxo7 knockout and control littermates were euthanized, different tissues were harvested, weighted and flash-frozen in liquid nitrogen until analysis. Tissues were lysed in RIPA buffer (50mM Tris pH 7.4, 150mM NaCl, 1% NP40, 0.1% SDS, 0.5% Sodium Deoxycholate, 1mM EDTA, in a 10:1 v\w ratio of lysis buffer to tissue sample) supplemented with phosphatase and protease inhibitors using a Dounce homogenizer. Lysates went through 3 cycles of freeze-thaw in liquid nitrogen followed by 15 minutes centrifugation at 14,000rpm at 4 C. Supernatants were collected, and protein concentration was calculated using the BCA protein assay. For electrophoresis, protein samples were loaded on a 4-12%, 15 well, Bis-Tris polyacrylamide gel. For Immuno-precipitation of flag-tagged PI31, spinal cords of Tg PI31:Chat^Cre^ and control littermates were harvested and flash-frozen in liquid nitrogen. Tissues were lysed in IP buffer (50mM Tris pH 7.4, 150mM NaCl, 0.5% NP40 in a 10:1 v\w ratio of lysis buffer to tissue sample) supplemented with phosphatase and protease inhibitors using a Dounce homogenizer. Lysates underwent 3 cycles of freeze-thaw in liquid nitrogen, followed by 15 minutes centrifugation at 14,000rpm at 4 C. Supernatants were collected and protein concentration was calculated using the BCA protein assay. Samples were adjusted to 5 mg/ml in a total volume of 600ul of IP buffer. 500ul of protein extract was incubated with 30ul of anti-Flag conjugated magnetic beads over night at 4 C on a rotator. Beads were washed 5x with wash buffer (20mM Tris pH 8.0, 100mM NaCl, 0.5% NP40, 1mM EDTA) and immune-precipitated proteins were eluted by incubating the beads in 50ul of 1x LDS sample buffer for 10 minutes at 90 C. For electrophoresis, protein samples were loaded on a 10%, 15 well, Bis-Tris polyacrylamide gel. Electrophoresis was carried out at RT (1 hour at 200V) in MOPS buffer (50 mM MOPS, 50 mM Tris Base, 0.1% SDS, 1 mM EDTA, pH 7.7). For immunoblotting, proteins in polyacrylamide gel were transferred to 0.45 mm PVDF membranes. Transfer was carried out at RT (90’, 100V) in Transfer Buffer (25mM Tris pH 8.3, 192M glycine, 20% Methanol). PVDF membranes were blocked in 5% BSA for 1 hour at room temperature and then incubated in primary antibody overnight at 4 C. Membrane washed 3x for 5 minutes in TBST (20mM TRIS pH7.4, 150mM NaCl, 0.1% Tween20) and incubated for 1 hour at room temperature with secondary antibody diluted in TBST. Membranes were washed 3x in TBST. Detection was performed with Amersham ECL Western Blotting Detection Reagent and imaged using ImageQuant LAS4000 (GE Healthcare) imager.

### Immunofluorescence analysis of muscle innervation

Whole mount Triangularis Sterni immunostaining protocol was adapted from Bril et. al.^26^. The thoracic wall containing the triangularis sterni muscle was dissected and fixed in 4% PFA for 1 hour on ice. The fixed tissue was then transferred to 0.25M glycine and the triangularis sterni muscle was dissected. The dissected muscle was incubated in permeabilization buffer (0.25M glycine, 0.5% TX-100 in PBS) for 30 minuets at RT and then transferred to blocking solution (0.25M glycine, 10% donkey serum, 0.5% TX-100 in PBS) over night at 4 C. After blocking, tissue was incubated with Alexa488 conjugated primary antibodies anti Tuj1 and anti Synapsin overnight at 4 C. To label post-synaptic acetylcholine receptors, samples were incubated in Alexa555 conjugated alpha-bungarotoxin for 30 minuets. The tissue was washed 3x for 30 minuets in PBS and mounted on a slide with Vectashield mounting media. Images were acquired with a Zeiss confocal LSM780 and Zen software.

### Image analysis

Image analysis methodology was performed as described previously^16^.

Quantification of axonal swellings: for the quantification of axonal swellings, z-stack images were taken at low magnification (10x). Maximal intensity Z-project images were generated using Fiji software and axonal swellings were counted manually. The average number of axonal swelling was calculated based on 3 images per animal. Image size – 850μm^2^.

Quantification of axonal sprouting: quantification of axonal sprouting was performed with the same images used for the quantification of axonal swelling using Fiji software tools. Briefly, for each image, 3 200×200px regions of interests (ROI) were chosen. Images were then skeletonized and the total axon length was measured for each.

### Statistical analysis

Welches t-test analysis (two tailed) were performed using GraphPad Prism version 8.4.1 for macOS, GraphPad Software, San Diego, California USA, www.graphpad.com

### Antibodies

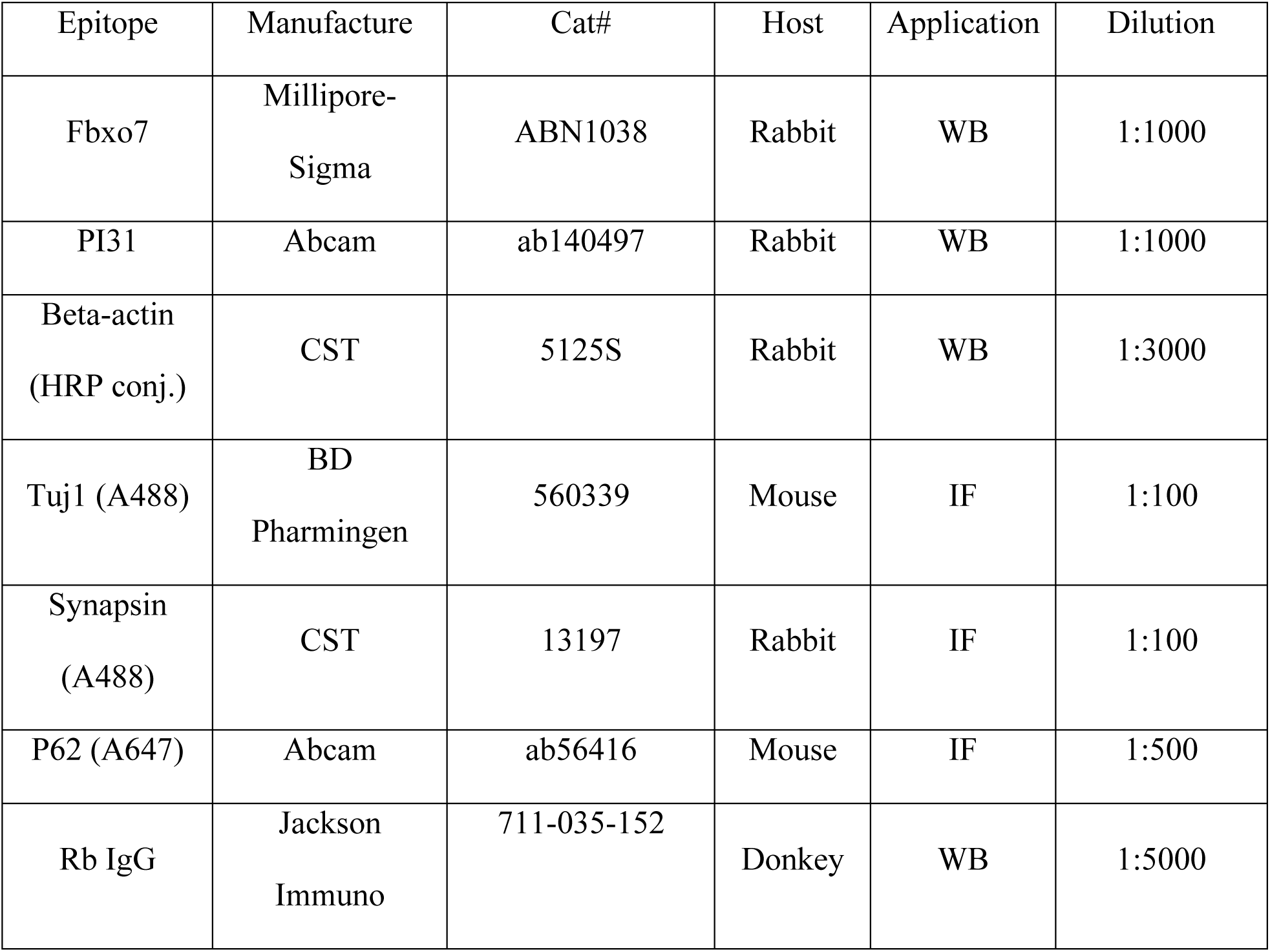

### Plasmids

pCAAGS\ES and pB-RAGE-GFP were a kind gift from Prof. Avraham Yaron, Weizmann Institute of Science, Rehovot, Israel.

### DNA oligos

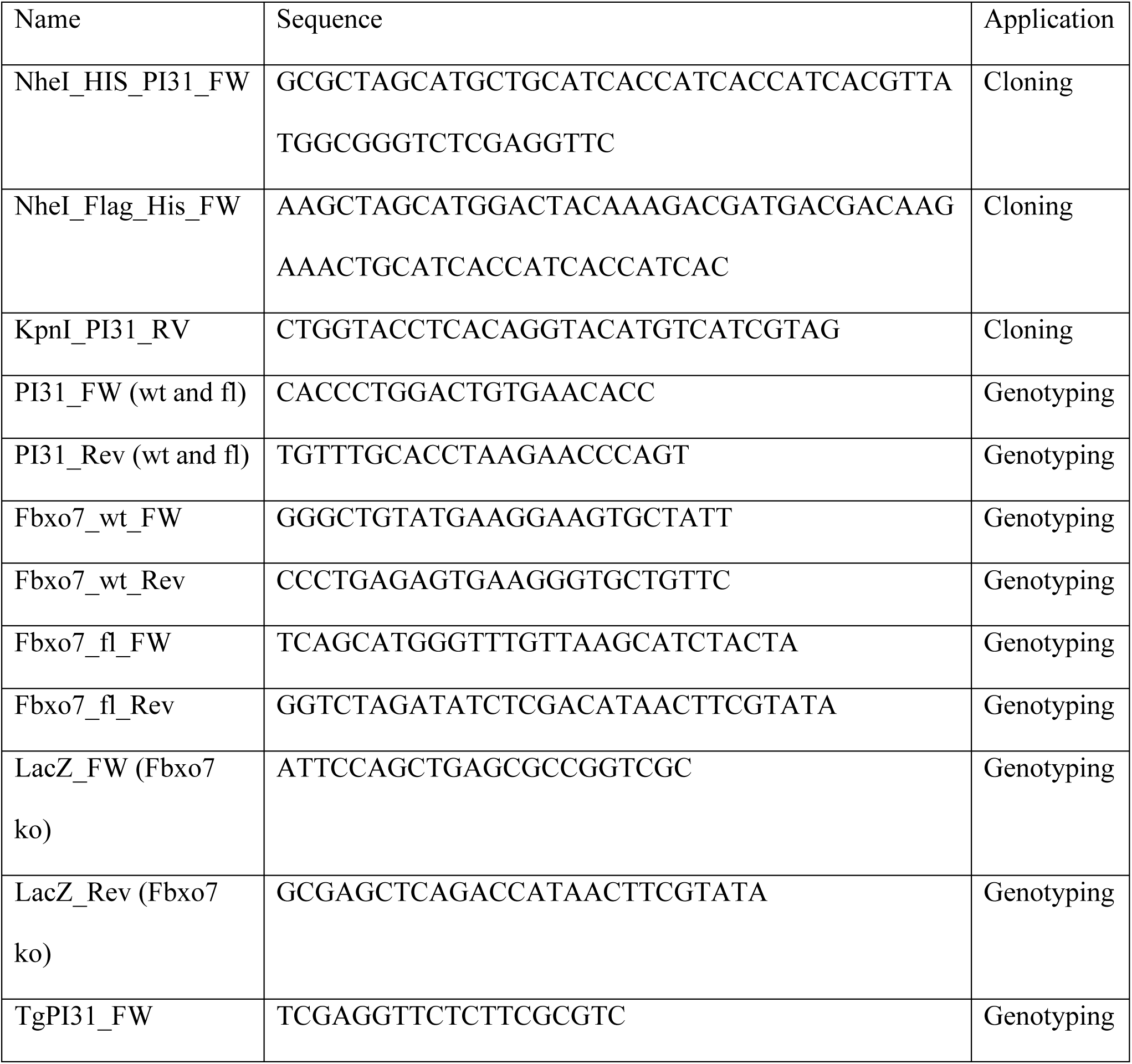
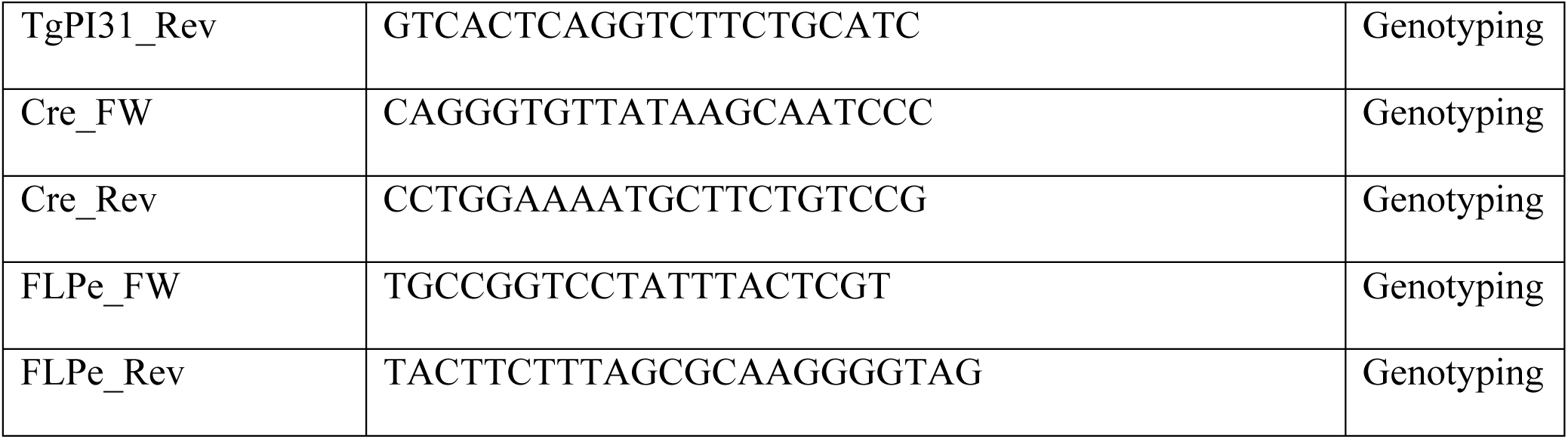

### Reagents

| Name | Manufacture | Cat# |
| --- | --- | --- |
| OneTaq Hot Start Quick-Load 2X Master Mix | New England Biolabs<br>(NEB) | M0488 |
| DirectPCR Lysis Reagent | Viagen Biotechnology | 102-T |
| Proteinase K, PCR Grade | Roche | 3115828 |
| SeaKem LE agarose | Thermo-Fisher Scientific | BMA-50004 |
| Tamoxifen | Sigma-Aldrich | T5648 |
| Corn Oil Delivery Vehicle | Sigma-Aldrich | C8267 |
| Phusion High Fidelity DNA Polymerase | Thermo-Fisher Scientific | F530 |
| FastDigest EcoRI | Fermentas | FD0275 |
| FastDigest NheI | Fermentas | FD0974 |
| FastDigest SalI | Fermentas | FD0644 |
| FastDigest Eco147I (StuI) | Fermentas | FD0424 |
| Tris Base Ultrapure | US Biological | T8600 |
| Sodium Chloride | Thermo-Fisher Scientific | S2713 |
| IGEPAL CA-630 (NP40) | Sigma-Aldrich | I8896 |
| Sodium Dodecyl Sulfate (SDS) | Thermo-Fisher Scientific | BP166-500 |
| Sodium deoxycholate | Sigma-Aldrich | D6750 |
| EDTA | Sigma-Aldrich | 03690 |
| MOPS | Sigma-Aldrich | M1254 |
| Glycine | Sigma-Aldrich | G7126 |
| Tween20 | Thermo-Fisher Scientific | BP337-500 |
| Halt™ Protease and Phosphatase Inhibitor | Thermo-Fisher Scientific | 78442 |
| Pierce BCA protein Assay Kit | Thermo-Fisher Scientific | 23227 |
| Anti-FLAG(R) M2 Magnetic Beads | Sigma-Aldrich | M8823 |
| Invitrogen Novex NuPAGE LDS Sample Buffer (4X) | Thermo-Fisher Scientific | NP0007 |
| NuPAGE™ 4-12% Bis-Tris Protein Gel | Invitrogen | NP0323BOX |
| NuPAGE™ 10% Bis-Tris Protein Gel | Invitrogen | NP0303BOX |
| Immobilon-P Membrane, PVDF | Sigma-Aldrich | IPVH85R |
| ALBUMIN FROM BOVINE SERUM | Sigma-Aldrich | A2153 |
| Amersham ECL Western Blotting Detection Reagent | Sigma-Aldrich | GERPN2134 |
| DPBS 10X | Thermo-Fisher Scientific | 14200075 |
| Paraformaldehyde 16% solution, EM grade | Electron Microscopy Sciences | 15710 |
| Triton X-100 | BIO-RAD | 161-0407 |
| Normal Donkey Serum | Jackson ImmunoResearch | 017-000-121 |
| Bungarotoxin A555 | Molecular Probes | B35451 |
| Vectashield | Vector Laboratories | H-1000 |

### Data Availability

No datasets were generated or analyzed during the current study.

## Acknowledgments

We would like to thank the transgenic and reproductive technology center of the Rockefeller University for their help in generating transgenic mouse strains, and members of the Steller lab, especially Dr. Jose A. Rodriguez, for thoughtful comments and helpful advice.

## Author contributions

AM and HS – conceptualization and methodology, AM – investigation and writing-original draft, AM and HS – writing-review and editing, HS – Funding acquisition and supervision.

## Extended Data

Extended Data includes:

Fig. S1

Captions for Movies S1 and S2

Other Supplementary Materials for this manuscript include the following:

Movies S1 and S2

## Supplementary Figures

**Figure S1.**
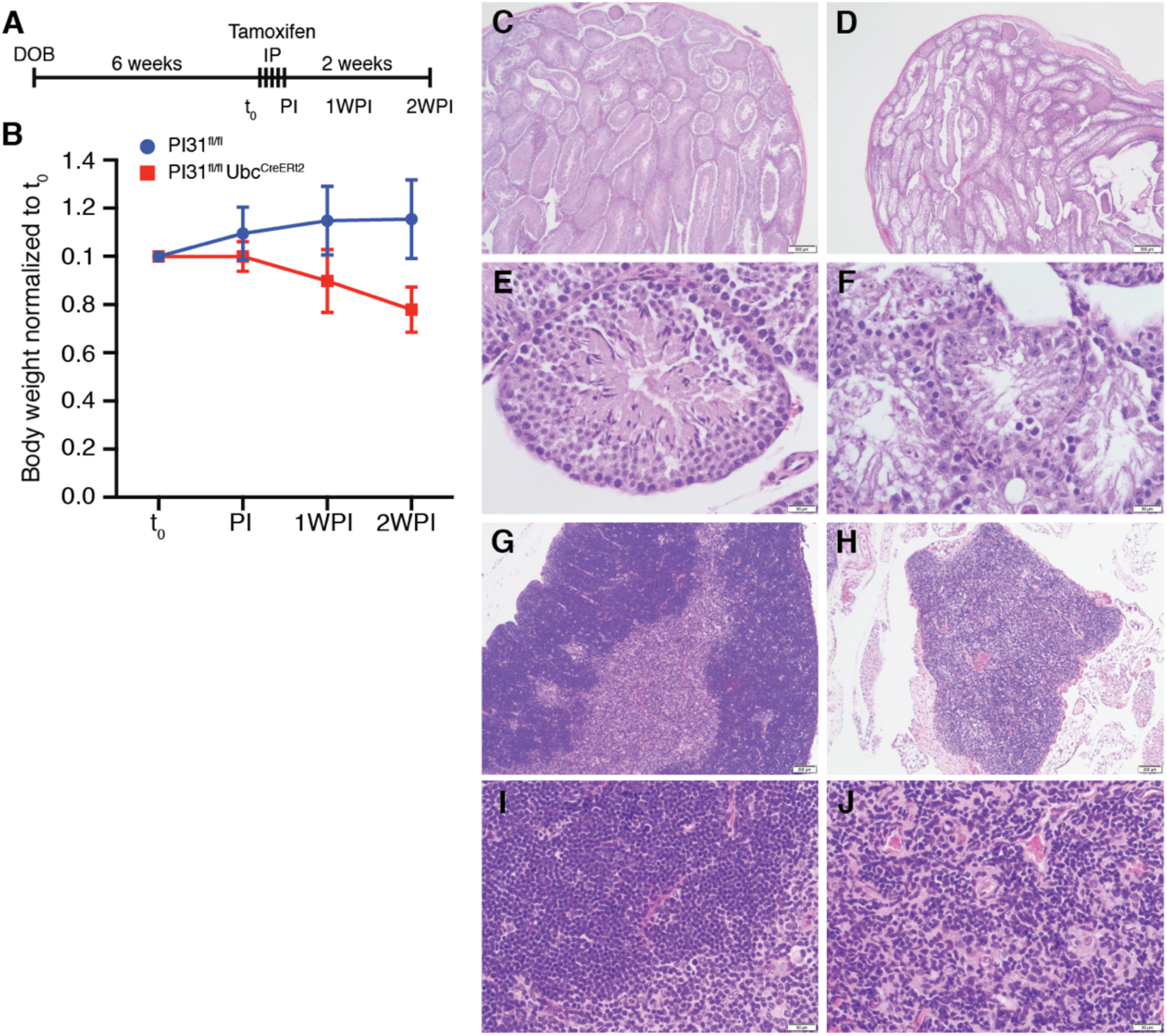
Inactivation of PI31 resembles loss of Fbxo7. (A) A diagram describing the tamoxifen-dependent induction of PI31 recombination in PI31^fl/fl^;Ubc^CreERt2^ mice. 6 weeks old mice were injected IP 100μl of tamoxifen at a concentration of 20mg/ml for 5 consecutive days. Mice were sacrificed for analysis 2.5 weeks after the last injection. (B) Normalized body weight of PI31^fl/fl^;Ubc^CreERt2^ and control littermates. Tamoxifen-injection resulted in weight loss in PI31^fl/fl^;Ubc^CreERt2^, but not in control littermates. (C-F) H&E stain of mouse testes of PI31^fl/fl^;Ubc^CreERt2^ shows a dramatic reduction in the number of mature spermatozoa, similar to what has been reported for Fbxo7 knockout mice^31^. (G-J) H&E stain of a mouse thymus of PI31^fl/fl^;Ubc^CreERt2^ shows a dramatic reduction in size, depletion of thymocytes, and loss of histological architecture.

**Movie S1.**

3-week old Fbxo7^fl/fl^::Chat^Cre^ mice were similar to control and Fbxo7^fl/fl^::TgPI31::Chat^Cre^ littermates. No significant size difference or motor dysfunction was detected. Displayed in the movie are two controls, two Fbxo7^fl/fl^;TgPI31;Chat^Cre^ animals, and one Fbxo7^fl/fl^;Chat^Cre^.

**Movie S2.**

8-week old Fbxo7^fl/fl^;Chat^Cre^ mice lost weight and exhibited motor dysfunction and breathing problems compared to control littermates. Fbxo7^fl/fl^;TgPI31;Chat^Cre^ mice were similar to control littermates, both in weight and motor function. Shown in the movie are the same mice that are displayed in Movie S1: two Controls, two Fbxo7^fl/fl^;TgPI31;Chat^Cre^ animals and one Fbxo7^fl/fl^;Chat^Cre^ mouse.

